# Improved sleep scoring in mice reveals human-like stages

**DOI:** 10.1101/489005

**Authors:** Marie Masako Lacroix, Gaetan de Lavilléon, Julie Lefort, Karim El Kanbi, Sophie Bagur, Samuel Laventure, Yves Dauvilliers, Christelle Peyron, Karim Benchenane

## Abstract

Rodents are the main animal model to study sleep. Yet, in spite of a large consensus on the distinction between rapid-eye-movements sleep (REM) and non-REM sleep (NREM) in both humans and rodent, there is still no equivalent in mice of the NREM subdivision classically described in humans.

Here we propose a classification of sleep stages in mice, inspired by human sleep scoring. By using chronic recordings in medial prefrontal cortex (mPFC) and hippocampus we can classify three NREM stages with a stage N1 devoid of any low oscillatory activity and N3 with a high density of delta waves. These stages displayed the same evolution observed in human during the whole sleep or within sleep cycles. Importantly, as in human, N1 in mice is the first stage observed at sleep onset and is increased after sleep fragmentation in Orexin-/- mice, a mouse model of narcolepsy.

We also show that these substages are associated to massive modification of neuronal activity. Moreover, considering these stages allows to predict mPFC neurons evolution of firing rates across sleep period. Notably, neurons preferentially active within N3 decreased their activity over sleep while the opposite is seen for those preferentially active in N1 and N2.

Overall this new approach shows the feasibility of NREM sleep sub-classification in rodents, and, in regard to the similarity between sleep in both species, will pave the way for further studies in sleep pathologies given the perturbation of specific sleep substages in human pathologies such as insomnia, somnambulism, night terrors, or fibromyalgia.

## Introduction

From the earliest investigations of brain electrical activity, researchers noticed signatures on the electroencephalogram (EEG) reflecting change in consciousness level in both humans (Berger, 1929) and animals (Caton, 1875). It became clear that mammalian sleep could be identified based on particular salient EEG events and the modification of oscillatory activity in larger frequency bands, which led to the subdivision of human sleep into five stages (Loomis *et al*., 1936). After the discovery of the rapid-eye-movements (REM) sleep (Dement and Kleitman, 1957), this categorization evolved slightly and converge towards the actual consensus with two major sleep states. First the REM sleep is characterized by desynchronized EEG, very low muscular tone, and rapid-eye-movements. Second, the non-REM (NREM) sleep presents irregular occurrence of various striking EEG events with a progressive apparition of spindles then slow waves with sleep deepening. Based on the latter events, NREM sleep was classified into four stages S1 to S4 (Rechtschaffen and Kales, 1968) and finally into the actual consensus of three NREM sleep stages N1 to N3 (Iber *et al*., 2007).

This type of classification is used in almost all sleep studies in humans, ranging from the physiological regulation of sleep to sleep pathology. Indeed, as pointed out by Hobson since the 60’s, “an important assumption of [these classifications] is that the variables to be measured by the method described are sensitive indices of biological events” (Hobson, 1969). The different processes occurring in NREM sleep are therefore likely to convey different functions (Kales and Kales, 1975). Accordingly, the N3 stage often called slow-wave sleep (SWS) due to the high prevalence of slow waves, is the deepest stage of sleep with the highest awakening threshold. It has been related to sleep pressure (Bersagliere and Achermann, 2010) and is homeostatically regulated (Achermann *et al*., 1993). This stage N3 corresponds to the periods with the highest release of growth hormone during sleep (Takahashi *et al*., 1968) and is associated with specific disorders such as night terrors and somnambulism (Kales *et al*., 1966; Fisher *et al*., 1973). On the other hand, N1 is the first stage to appear at sleep onset and within sleep cycles, and has the lowest awakening threshold (Emmons and Simon, 1956; Roth, 1961). It can be associated with hypnagogic imagery (Dement and Kleitman, 1957; Carskadon and Dement, 2011) and is strongly upregulated by sleep fragmentation observed in several pathological situations (Levine *et al*., 1987). This stage is for instance increased in fibromyalgia (Wu *et al*., 2017), narcolepsy with cataplexy (Dauvilliers *et al*., 2003, 2007) and in persistent primary insomnia (Merica and Gaillard, 1992). Interestingly, fatal familial insomnia is a genetic sleep disorder associated with thalamic hypometabolism and a massive decrease in SWS whereas N1 duration is preserved, suggesting that these two phases of NREM sleep are differently regulated (Montagna, 2005). Moreover, it has also been shown that the increase in transitions from N2 to N1 is a better predictor of insomnia disorder that any conventional sleep parameters (Wei *et al*., 2017).

Despite their clinical relevance, the different stages of NREM sleep have never been considered in experimental research on rodents. In pharmacological studies designed to characterize the efficiency of drugs on sleep, results in human showed that drugs affect the different substages of sleep in a specific manner but fail to provide the same specificity in rodent, and only address global measure of sleep (Brisbare-Roch *et al*., 2007; Gotter *et al*., 2016). The main limit is the rather vague or inappropriate definition of the NREM sleep in rodent that precludes efficient translational study, even though the pharmacology of vigilance states perturbations is very similar between those species (Toth and Bhargava, 2013). Rodent NREM sleep is often considered by default as a uniform state and most scientific articles use the terms “SWS” and “NREM” indifferently, which may lead to misinterpretation when comparing human and rodent literature. Moreover, when researchers attempt to perform a direct comparison of sleep properties across species, rodents sleep is compared to the sole human stage 2 of NREM sleep (Mölle *et al*., 2009, section Materials and methods), making difficult any further comparison in sleep regulation.

In addition to a substantial improvement of translational studies on sleep physiopathology, the field of memory research is certainly the most likely to benefit from taking into consideration the different substages of NREM sleep. Indeed, very fine neurophysiological mechanisms related to memory consolidation were conserved in rodent and human (Rasch and Born, 2013). To date, two main hypotheses have been proposed in order to explain the benefit of sleep on memory functions. On one hand, the synaptic homeostasis hypothesis (SHY) posits that sleep serves a global decrease of synaptic weights allowing restoration of the ability for further learning (Tononi and Cirelli, 2003, 2014a). This synaptic downscaling would rely on slow waves, which constitute both a marker of sleep pressure and an actor in the homeostatic process as it is thought to induce long-term depression (LTD) during sleep. On the other hand, accumulation of evidence supports the two-stage consolidation process model, in which hippocampal sharp wave ripples related neuronal reactivations during sleep allow the active consolidation of memory traces and the transfer of information from hippocampus to prefrontal cortex for long-term storage (Buzsáki, 1989; Wilson and McNaughton, 1994; Lee and Wilson, 2002). Those two hypotheses may appear incompatible and are the subject of an intense debate. However, Genzel and colleagues (Genzel *et al*., 2014) recently suggested that distinguishing two NREM substages, a light and a deep sleep, could partly explain those discrepancies and reconcile the two theories: consolidation during light sleep and down-scaling during deep sleep. However, this remains speculative since there is still no clear definition of what is light versus deep sleep in rodents.

Altogether, the lack of similar methodology to describe sleep in humans and rodents remains a critical drawback. The convergence in sleep scoring methodology across species is thus critical to favor translational research and the extrapolation of animal findings to humans. In the present article, we propose a more precise and comprehensive sleep scoring method in rodents that considers the heterogeneity of NREM sleep and that is inspired by sleep scoring in humans in order to encourage translational studies on sleep.

## Materials and Methods

### Experimental Design

Controlled laboratory experiment were conducted on a total of 25 C57Bl6 male mice (Mus musculus), 3–6 months old, and 3 Orexin knock-out mice (Orexin^-/-^). Polysomnographic recordings from 20 healthy subjects free from medication were obtained from the DREAMS database (see more details below). No blind experiment or subject randomization were used.

### Surgical protocols and behavior experiments on mice

All mice underwent stereotaxic microsurgery for electrode implantation. Simple tungsten wires were lowered in the prelimbic cortex (AP +1.9, ML 0.4, DV −1.6) and in the CA1 pyramidal layer of hippocampus (AP +2.2, ML +2.0, DV −1.0) of the right hemisphere. Among those, 15 mice were also implanted with tetrodes in the prelimbic cortex to record single-unit activity, and 10 mice with tungsten wire in the neck muscles to record EMG activity. During recovery from surgery (>3d) and during all experiments, mice were housed in an animal facility (12h light/12h dark, constant light and monitored temperature), one per cage, and received food and water ad libitum. All mice were free of any manipulation before being included in this study. Natural sleep in mice home cage was continuously recorded during the light phase (9am to 9pm).

All behavioral experiments were performed in accordance with the official European guidelines for the care and use of laboratory animals (86/609/EEC) and in accordance with the Policies of the French Committee of Ethics (Decrees n° 87–848 and n° 2001–464). Animal housing facility of the laboratory where experiments were made is fully accredited by the French Direction of Veterinary Services (B-75-05-24, 18 May 2010). Animal surgeries and experimentations were authorized by the French Direction of Veterinary Services for K.B. (14–43).

### Electrophysiological recordings and analysis

Signal was sampled at 20 kHz, digitalized and amplified by INTAN system (RHD2000-series). Local field potentials were sampled and stored at 1.25 kHz. Recordings were band-pass filtered between 0.6–9kHz, processed using NeuroScope (Hazan *et al*., 2006), and single-units were sorted using KlustaKwik (Harris *et al*., 2000). Classification between interneurons and pyramidal neurons was based on firing rate, autocorrelogram and spike waveform (Sirota *et al*., 2008). See supplementary Methods for further details.

### Classical sleep scoring

To determine sleep periods, a 30Hz camera placed 1m above the cage videotaped the mouse and a Matlab-based program (MathWorks, Natick, MA) analyzed frame-by-frame image difference to determine sensitively the movements of the animal. Immobility threshold was set above respiratory movements. For a majority of recordings, movements from video were replaced by accelerometer signal from the INTAN headstage, without changing our ability to differentiate sleep and wakefulness. REM and NREM sleep were distinguished by automatic k-means clustering of the theta/delta ratio extracted from the power spectrograms of hippocampal LFP signal during the episodes where the animal was immobile. REM sleep corresponded to high theta/delta ratio periods, NREM sleep to the rest of sleep. Importantly, we did not further differentiate “intermediate sleep” or “transitory sleep”, classically defined as a high theta/delta ratio in the hippocampus but remaining spindles and delta activity in the prefrontal cortex occurring at the transition from NREM sleep to REM sleep sleep (Gottesmann, 1996; Datta and Hobson, 2000). Applying our classification, this intermediate sleep is included in the REM sleep sleep, due to the high theta/delta ratio, and cannot account for the results on NREM substages.

### NREM sleep rhythm detection

Offline delta wave detection used two LFP channels recorded in the prefrontal cortex, ideally one in superficial layer and the other in deep layer. In the absence of the latter configuration, two channels at different depth were selected so that the dipole generated by delta waves was clearly apparent. Those two signals were band-pass filtered (1-20Hz) and subtracted to obtain the differential signal between depth and superficial layers. Delta waves were then defined as extrema of this filtered differential signal in the 1-5Hz frequency band, above a threshold of 2 standard deviations (sd) defined during sleep and that last at least 75ms.

For offline detection of “oscillating periods”, LFP signal at 1250Hz from prefrontal cortex is filtered between 2 to 20Hz, by 2Hz steps (2-4Hz, 4-6Hz,…). For each frequency range, sd is calculated and signals above 2sd are detected as oscillating events. All events are pooled, and period corresponding to burst of events (at least 3 events with an inter-event period smaller than 1s) are determined as oscillating periods.

### NREM sleep classification method

As summed up in Supplementary Fig.1, scoring of Wake, REM and NREM sleep was classically defined. Movement episodes shorter than 3s was considered as sleep, and immobility episode shorter than 3s as wakefulness. Two subsequent episodes scored as REM sleep and separated by less than 5s were merged into one single REM sleep episode. Episodes scored as REM or NREM sleep shorter than 3s were dropped and scored as the previous stage. Then, on NREM sleep periods we automatically detected “oscillating periods” in the low frequency band (2-20Hz) including delta oscillation and spindles to distinguish between non-oscillating period N1 and oscillating periods N2/N3. Two episodes scored as N1 and separated by less than 3s were merged into a unique N1 episode, and N1 episode shorter than 1s was dropped and scored as the previous stage. Delta wave bursts periods were then detected on N2/N3 epochs with inter-delta intervals <700ms and scored as N3. N3 episodes separated by less than 3s were merged as one N3 episode. The remaining N2/N3 epochs was scored as N2, and N2 episodes shorter than 1s was dropped and scored as the previous stage. Change in the criteria of inter-delta intervals to determine delta bursts (<1s instead of <700ms), did not modify the result of NREM substages classification as percentage of each substages was conserved.

### Human recordings

Polysomnographic recordings from 20 healthy subjects free from medication (16 women and 4 men, from 20 to 65 years old, mean 33,5) were obtained from the DREAMS database (2013, http://www.tcts.fpms.ac.be/~devuyst/Databases/DatabaseSubjects/) of MONS University-TCTS Laboratory, and Brussels University-CHU de Charleroi-Sleep Laboratory. Sleep scoring was performed by an expert from the Sleep Laboratory, by 30s epochs according to criterions from the AASM20 (Iber *et al*., 2007). Briefly, sleep begins when both muscular tone and occipital alpha band decrease. The latter epoch and the following ones are scored as N1 unless spindles or K-complex appear in the EEG, in which case the epoch is scored as N2. Unless arousal or rapid eye movements occur, following epochs are scored N2 until slow oscillations get frequent and last at least 20% of the epoch (cumulative). This epoch is then scored N3, a stage also called slow wave sleep or deep sleep (Iber *et al*., 2007). Information on sleep sessions is summarized in Supplementary Table.1)

## Results

To characterize the sub-structure of NREM sleep in mice, we performed extracellular electrophysiological recordings during natural sleep in 25 mice (N=25; total sleep sessions: n=65) implanted with single wires for local field potentials (LFP) recordings in the pyramidal layer of the hippocampus and in superficial and deep layers of the medial prefrontal cortex (mPFC). Fifteen of these mice were also implanted with tetrodes in the mPFC to record spiking activity. Rodent sleep characteristics were compared to human sleep from the DREAMS database consisting of polysomnographic sleep sessions from 20 subjects scored according to AASM on epochs lasting 30s (See Methods, Supplementary Table.1, (Iber *et al*., 2007)).

### NREM substages definition in mice and comparison with human sleep

LFP recorded in mPFC and hippocampus during mouse natural sleep allow to dissociate between two classical sleep stages: REM sleep during which the hippocampus oscillates at theta frequency (6-8Hz), and NREM sleep associated with a range of various cortical rhythms such as spindles and delta waves. A closer look during NREM sleep reveals an important heterogeneity mostly related to the density of delta waves (Fig.1A) corresponding to sleep periods ranging from no delta waves and no spindles, irregular occurrence of delta waves, to periods with strong and constant bursts of delta waves.

**Fig. 1.**
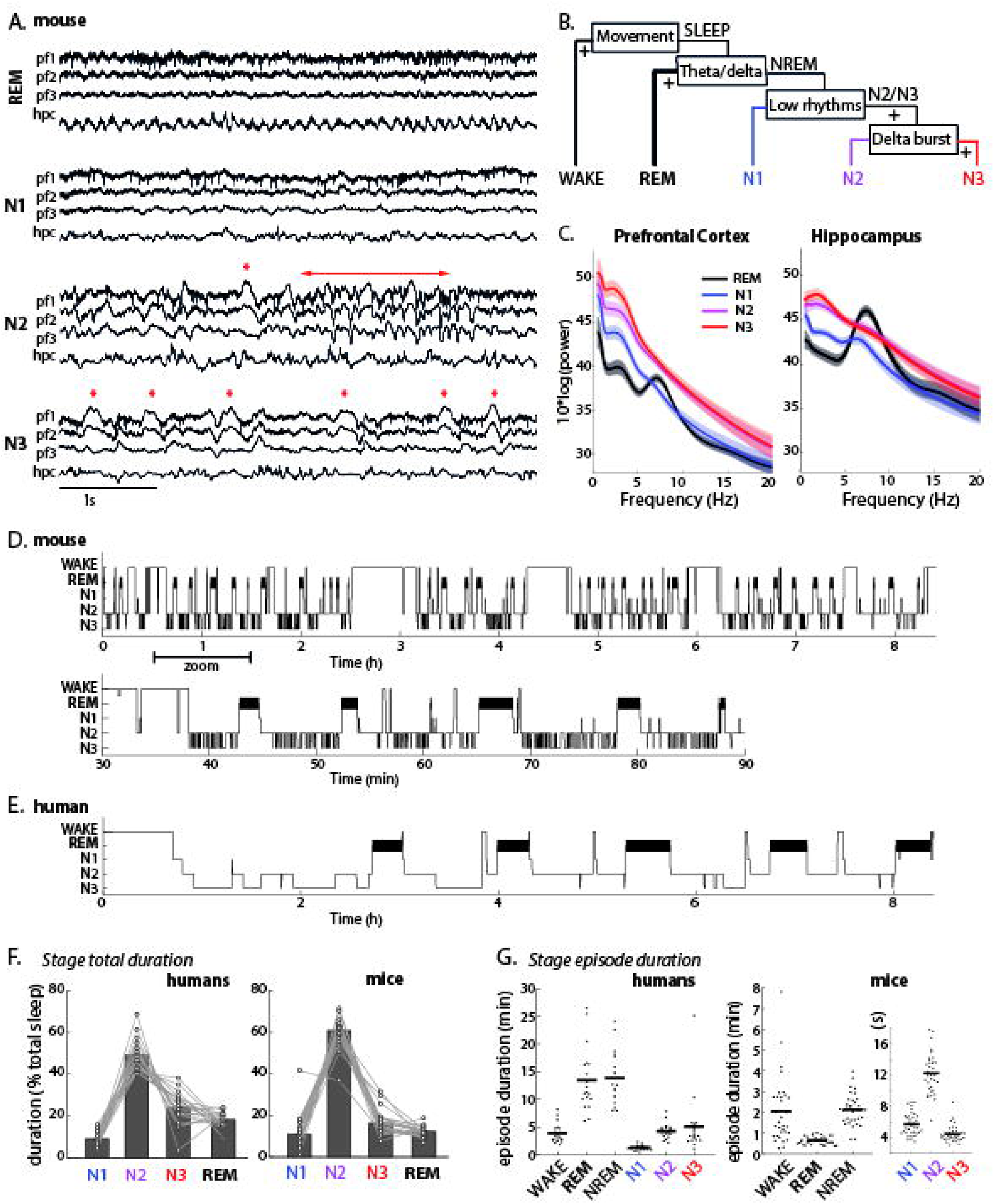
NREM sleep scoring in mice, and comparison between human and mouse sleep characteristics: hypnograms and stage durations. **(A)** Example of electrophysiological signals recorded in prefrontal cortex and hippocampus at different times of the same sleep period pfc: prelimbic prefrontal cortex, hpc: dorsal hippocampus. Asterisks and arrow indicate prefrontal delta wave and spindle, respectively. **(B)** Diagram of sleep scoring distinguishing different NREM stages based on low frequency oscillations. See Methods for further details. **(C)** Power spectrum recorded in pfc (left, N=15 mice) and hpc (right, N=14 mice) averaged during sleep stages. Note the gradual increase of delta frequency band (2-4Hz) between N1, N2 and N3; and the theta band (7-10Hz) characteristic of REM sleep. **(D)** Example of mouse hypnogram over the whole sleep session or with an increased time resolution (zoom in, bottom hypnogram). Note the natural fragmentation of mouse sleep with long periods of wakefulness. **(E)** Example of human hypnogram across one night of sleep shows similar architecture in terms of regular REM although the time scale is ten times larger (30s scoring epochs according to AASM, DREAMS database). **(F)** Total amount of sleep substages as a percentage of total sleep in mice (left, N=25 mice, averaged on 2–4 nights) and human subjects (right, N=20). **(G)-** Averaged episode duration for all 5 vigilance states. Each dot represents one recording session; bar represents the mean.

Given the importance of delta waves in this classification, we decided to optimize the method of delta waves detection. Delta waves are visible in the LFP as a negative deflection in superficial layers and a positive deflection in deep layers, characteristic of an electrical dipole, and are associated with silence of the neuronal spiking activity (down states). Prefrontal cortex LFP was thus recorded at two different depths in order to identify with high certainty true delta waves (Calvet *et al*., 1973), identified as an inversion between signals from the two electrodes (Fig.1A, and Methods). This prevented false positives due to random fluctuations in the low frequency band.

The sleep scoring steps are described in Fig.1B (and further detailed in Methods and Supplementary Fig.1). First, we automatically scored sleep and wakefulness based on movement detection. Similar results were obtained using either video or accelerometer (n=65/N=25), or electromyogram (n=29/N=10). We then discriminated NREM and REM sleep using theta/delta ratio from hippocampal LFP (see Methods). During the NREM sleep epochs, periods with no oscillations in the low frequency band (2-20Hz) including delta oscillation and spindles were classified as N1, while periods with bursts of delta waves (see Methods) were scored as N3. The remaining NREM sleep epochs were scored as N2. It is worth mentioning that this scoring method is continuous on the contrary to most classical sleep scoring performed on pre-determined times windows in both human and rodents.

Changes in the oscillatory activity were visible in the averaged power spectrum of LFP computed in the different sleep stages (Fig.1C): theta oscillations were strong during REM sleep in the hippocampus and power in the low frequency band (2-4Hz) in mPFC gradually increased from N1 to N3. Interestingly, these modifications are clearly visible when looking at the transitions between states (Supplementary Fig.2). As expected, mPFC power drops at transition to wakefulness, theta band power in hippocampus shows a strong increase at transition to REM sleep, and a drop of movement and EMG are seen at transition to sleep. This shows that our automatic sleep scoring, before any NREM subdivision, gives similar results than classical sleep scoring in mice.

We then computed hypnograms representing mouse sleep evolution across time. At first sight, the hypnograms share strong similarities with human sleep (Fig.1D-E) but mice sleep seems to be more fragmented, interrupted by long wake episodes, and with more frequent transitions with sleep substages (Fig.1D).

In mice, N1, N2, N3 and REM sleep represented respectively 10.0+/-0.9, 61.4+/-0.9, 16.2+/-0.8, 12.3+/-0.5 percent of total sleep, similar to the global fractions of substages in human sleep: 9.1+/-0.8, 49.0+/-1.7, 23.8+/-1.7 and 18.2+/-0.8 respectively (Fig.1F). The proportion of the different NREM substages is highly reproducible across mice. Moreover, the individual variability in the substages repartition from one mouse sleep session to another, was smaller than the inter-individual variability for N1 (inter-variability 0.73 versus intra-variability 0.32 +/-0.21), N2 (0.11 vs 0.08 +/-0.05) and N3 (0.35 vs 0.20 +/-0.19), emphasizing that each mouse has its own sleep signature similar to what observed in humans (De Gennaro *et al*., 2008; Gander *et al*., 2010). The main difference between mice and human sleep stages, and especially the NREM substages, concerns the average episode duration. Individual episodes last a few seconds in mice, compared to minutes in humans (Fig.1G). This brevity may account for the neglect of NREM substages in the rodent literature but our analysis reveals that despite the different time scales, most properties are shared between the two species.

### Similar dynamic of NREM substages across the night in mice and humans

We first compared the dynamics of NREM substages in human and rodent across the whole sleep period, starting at the beginning of bedtime for humans and of the light phase for mice (Fig.2). As expected REM sleep tend to increase with time during sleep and N3, defined as a state with high density of delta, was more present at the beginning of sleep and its occurrence decreased with time, consistent with the homeostatic regulation of slow waves (Borbely, 1982). Importantly, the shape of the delta wave events is almost identical across the whole sleep session excluding that the evolution is biased by a problem of detection (Supplementary Fig.3C, less that 5% modification of amplitude).

**Fig. 2.**
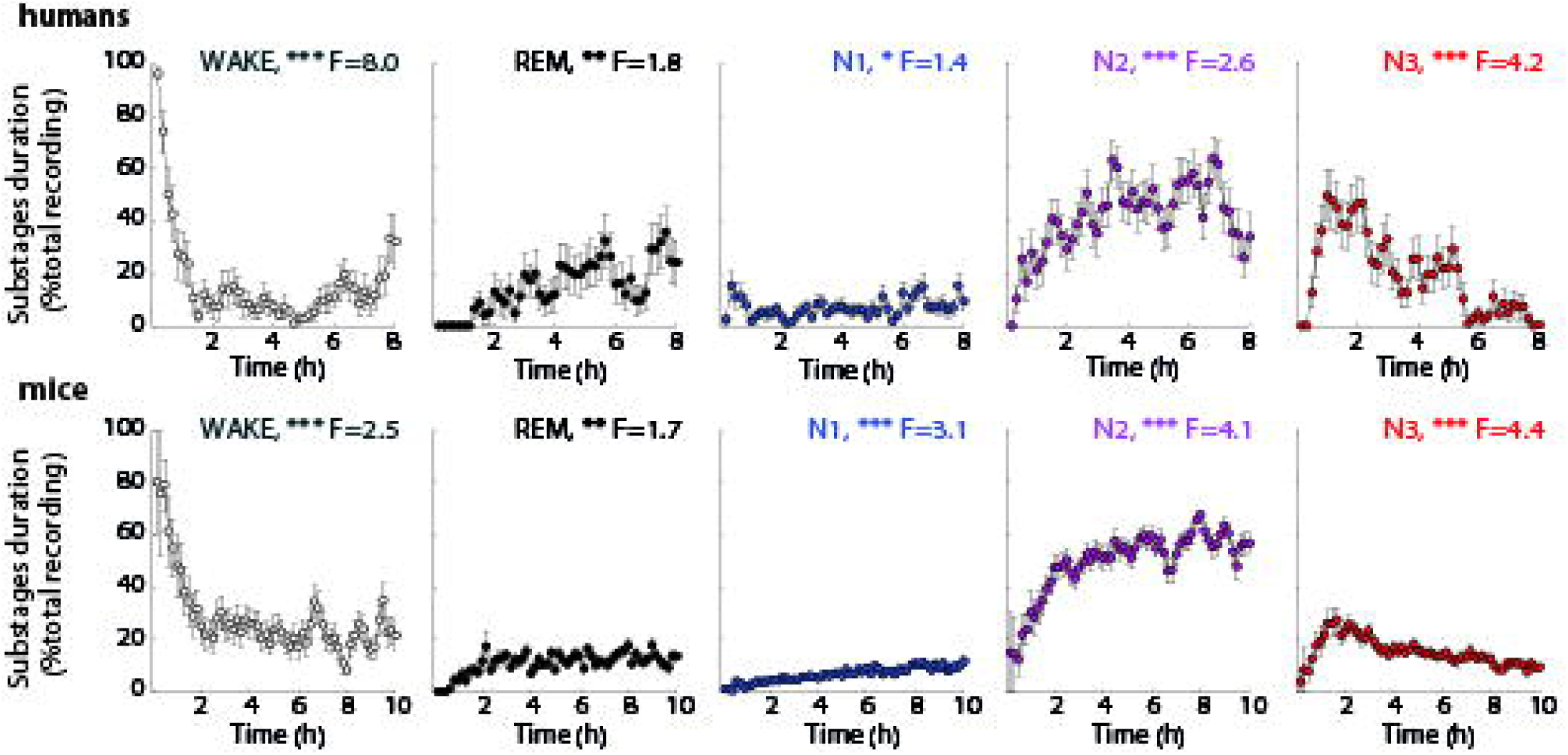
Mice and human’s NREM stages similarity at whole night time scales. Evolution across sleep period of stages duration, comparison between humans (upper panel, N=20 subjects) and mice (bottom panel, n=65 sessions, N=25 mice). Time zero corresponds to recording start for humans and light onset for mice. ANOVA reveals a significant effect of time for all stages; N1, N2 and REM amount tend to increase over sleep period, while N3 decreases with time.

More interestingly, the two other NREM substages N1 and N2 also showed the same evolution across sleep in both species. This validates our strategy to consider these sleep epochs as individual NREM substages and suggests that the global organization of sleep states follow the same global evolution in human and mice.

### Similar dynamic of NREM substages within sleep cycles in mice and humans

One strong property of REM sleep is to occur periodically during sleep, which led to the definition of sleep cycles, classically defined as the period between the end of two successive REM sleep episodes, excluding long periods of wake. Visual inspection of mouse hypnogram (Fig.1D, zoom in) shows that REM sleep episodes are indeed regularly spaced. Accordingly, the distribution of sleep cycle duration shows a clear pic both in human and rodent (Fig.3A). As previously shown, the mean duration of a sleep cycle is around 10 minutes in rodent and 90 minutes in human (Dement and Kleitman, 1957; Trachsel *et al*., 1991).

**Fig. 3.**
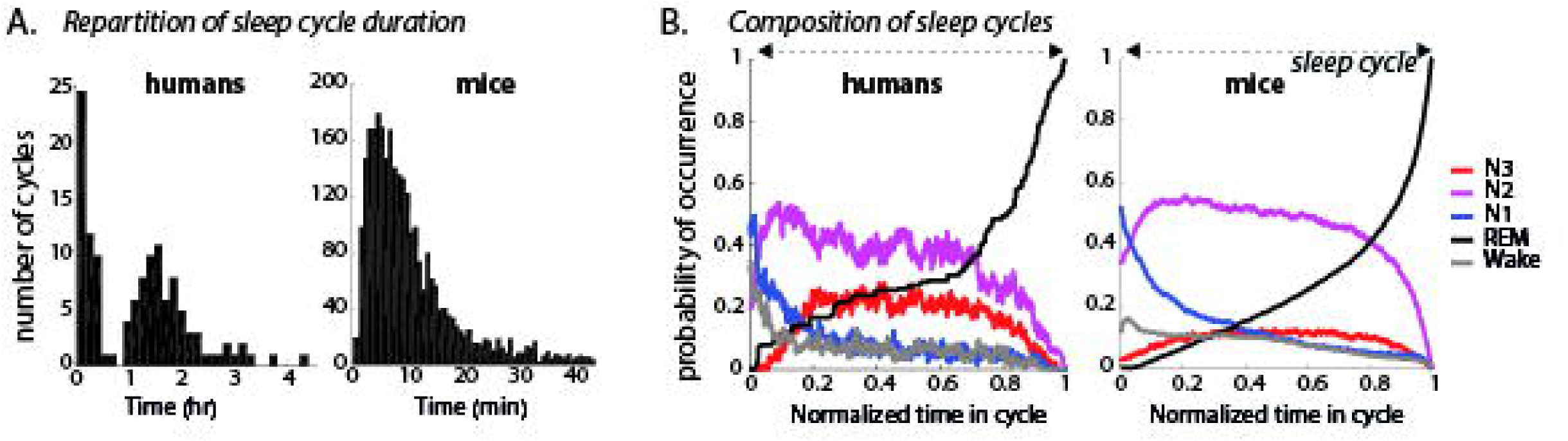
Mice and human’s NREM stages similarity in sleep cycles. **(A)** Distribution of sleep cycle durations for humans (N=20) and mice (n=65 sleep sessions, N=25 mice). One cycle is defined as the period between the end of two successive REM episodes. **(B)** Stages distribution within sleep cycle, comparison between humans and mice. Averaged probability of occurrence for each stage along the advancement of the cycle, normalized between 0 (beginning of the cycle) and 1 (end of the cycle) for all subjects (left, N=20) and all mice (right, N=25).

Importantly, the similarity of sleep substages between mice and humans is also visible at the scale of sleep cycles. When averaging the content of all sleep cycles, a striking similarity between the respective sleep stages occurrence in mice and humans can be observed (Fig.3B). Sleep cycles in mice and humans are, on average, revealing the same structure: cycles mostly start by wakefulness and N1 stage, then N2 stage, followed by N3, up to the apparition of REM sleep that terminates the cycle. In addition, N3 sleep stage is overrepresented in the first sleep cycles, whereas the last cycles are composed of more REM sleep (Supplementary Fig.4).

Altogether, these results show that the dynamics of sleep in mice are very similar to what is observed in humans at both time scales: sleep period and sleep cycles.

### Short scale dynamics of NREM sleep in mice and human

To further understand the dynamics of NREM microstructure, we analyzed the transitions between the sleep substages (Fig.4) As expected, the number of transitions is higher in mice compared to humans, due to the short duration of substages (Fig.1G). However, the repartition of transitions is highly conserved in the two species as evidenced by diagrams in which both the global durations of stages and the number of transitions are represented (Fig.4A, see also Supplementary Table.2). The most frequent transitions are N2-N3-N2 and N3-N2-N3 in both species showing the imbrication of these two stages (Fig.4B). Similarly, there is a high prevalence of transitions considered as a deepening of sleep (e.g., N1-N2-N3 or Wake-N1-N2). The similarities between mice in human are further confirmed by the significant correlation between the frequencies of transitions between humans and mice (Fig.4C). Altogether, this shows that the microstructure of NREM is conserved from human to rodent at large scale, at the scale of a sleep cycle but also when considering the sequences between the different substages.

**Fig. 4.**
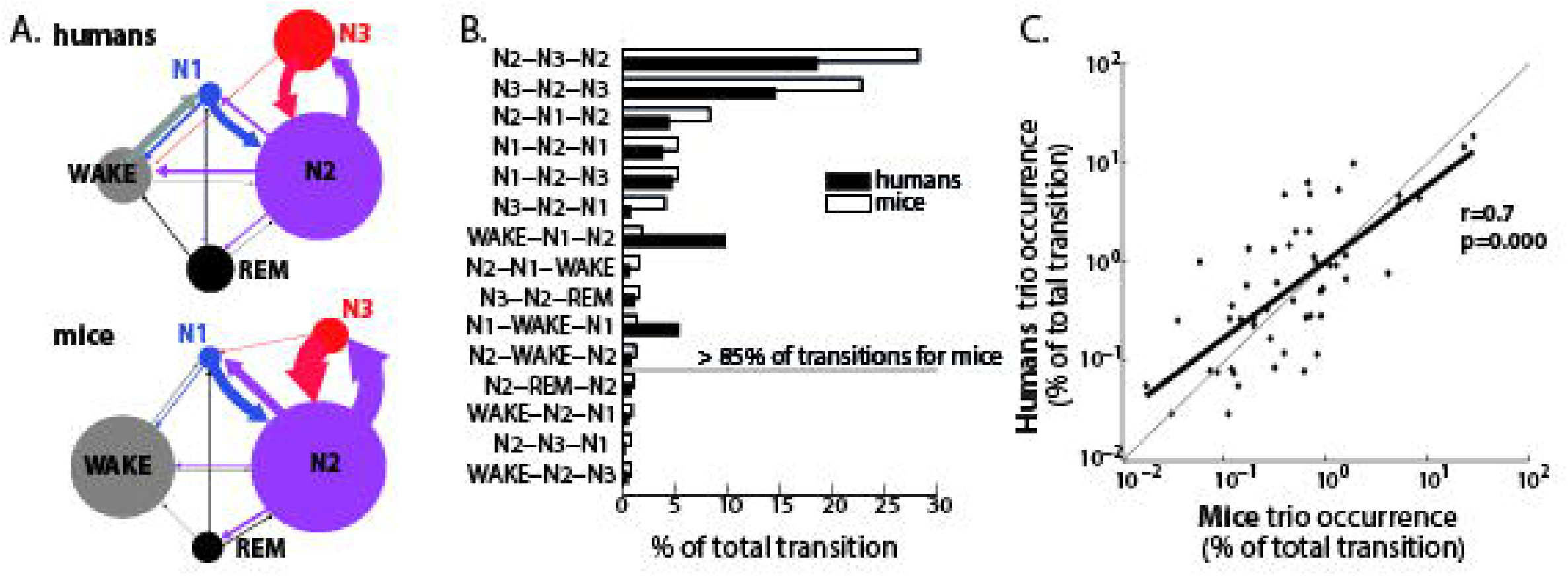
Mice and human’s transition between NREM stages. **(A)-** Diagram of transition between stages. Circle size is proportional to total amount of each stage, arrows thickness to the number of transition from one stage to another. Note the frequent switch between N2 and N3 in mice, and the frequent N2-N1 transition which is almost absent in human probably due to continuity rule in sleep scoring (after N2 period, score N2 even in the absence of K-complex or spindles until other stage characteristics occur, see (Iber *et al*., 2007)). **(B)-** Repartition of 3 stages sequence transitions ordered by frequency of occurrence in mice sleep. Only 15 out of 80 possible transitions are shown. Note that the most frequent ones are the same for mice and humans. **(C)-** Correlation between percentage of all 80 possible trio of transitions. Note that most frequent ones are around 30% (see B.) and the less frequent ones are below 0.05%, for both humans and mice.

### N1 sleep in mice is distinct from wakefulness or other NREM substages, and shares common properties with human stage 1

A crucial observation is the presence in mice of a stage similar to the human stage 1. Several reports have shown that sleep periods with low amplitude LFP can be observed in rodent. Here we show that these periods are not randomly observed across sleep but are in fact reminiscent of the N1 stage observed in human.

First, we confirmed by using electromyogram and the quantity of movement, that N1 was not associated to any increase of movement, unlike the transition to wakefulness (Supplementary Fig.5). We performed additional control by using a new method our Lab recently published, in which we showed that the gamma activity in the olfactory bulb allows to differentiate wakefulness from sleep without the need to consider motor activity to define sleep (Bagur *et al*., n.d.). Consistent with our hypothesis, gamma activity in the olfactory bulb is almost absent during N1 and does not show any modification at transition toward N1 contrary to what observed at transition to wakefulness.

Interestingly, transitions to N1 - defined with LFP signal - are also associated to a massive modification of neuronal activity. We recorded 591 singles units (N=15 mice) in mPFC, and analysis of the firing rate at transition to N1 reveals a sharp modification of neuronal activity with some neurons being increased while other are decreased (Fig.5A). Importantly, the sharp deflections of neuronal activity are larger when considering transitions to N1 that transitions to other substages (Supplementary Fig.6).

**Fig. 5.**
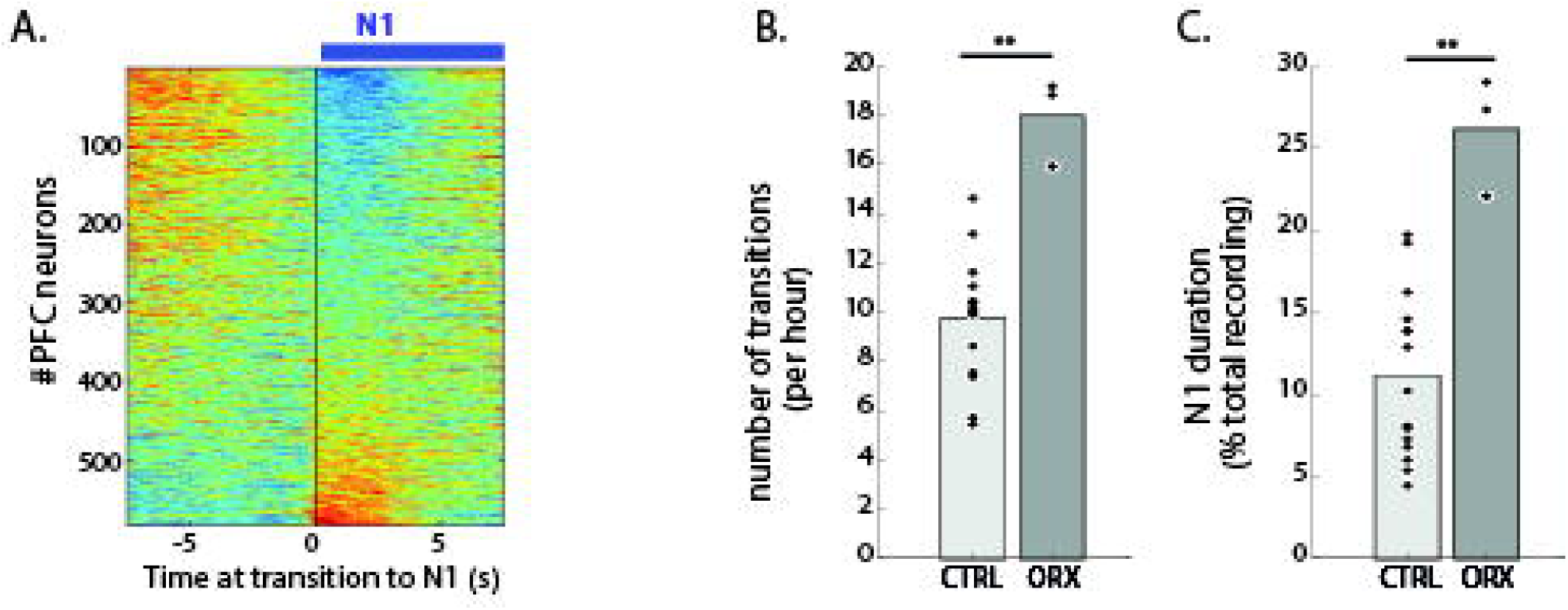
N1, a particular state associated to fragmented sleep in Orexin-/-mice. **(A)** Peri-event histogram of prefrontal cortex neurons (n=591 neurons) activity at transition to N1, ordered by the intensity of response to transition. Each row corresponds to the cross-correlogram of one neuron spikes by all transition to N1, normalized by zscore. **(B)-** Number of transition from sleep to wakefulness in Orexin-/-mice (ORX) and controls (CTRL) in the light phase of the day. Note the increased fragmented sleep in Orexin-/-mice (Wilcoxon: p=0.025). **(C)-** Time spent in light sleep N1 expressed as a percentage of total sleep in Orexin-/- mice (ORX) and controls (CTRL) (Wilcoxon: p=0.025).

This further support that N1 in mice is indeed a state of sleep with specific LFP and neuronal properties that shares homology with human stage 1 regarding its evolution across the whole sleep session or within sleep cycles. In order to provide additional evidence of the similarity of N1 stage in both species, we investigate whether N1 in mice sleep displays the same susceptibility to sleep fragmentation to the one observed in human. Indeed, sleep fragmentation in humans, observed in pathological situation such as narcolepsies or insomnia, is usually associated with an increase in N1 sleep (Dauvilliers *et al*., 2003, 2007; American Psychiatric Association, 2013). We thus tested whether N1 duration in mice showed a similar increase along with pathological sleep fragmentation. We used Orexin knock-out mice, a well-described model of narcolepsy, that have been shown to display an increased in sleep fragmentation without any change to the total amount of sleep or wakefulness (Chemelli *et al*., 1999; Willie *et al*., 2003; Diniz Behn *et al*., 2010). We thus recorded natural sleep in Orexin-/- mice and confirmed a higher fragmentation of sleep compared to controls (Fig.5B-C), with no change in the global percentage of wakefulness (Control: 25.7%+/-7.0; Orexin-/-: 27.9%+/-13.6). Importantly, this increase in sleep fragmentation was associated with an increase of N1, demonstrating that N1 in mice shares the same properties than human N1.

### NREM substages and the dichotomy between light and deep sleep in synaptic downscaling

Two main theories have been proposed to explain the beneficial role of sleep on memory that have been opposed regarding the type of plasticity required. The reactivation/consolidation theory (Buzsáki, 1989) posits that synapses important for memory are strengthened during sleep, while SHY (Tononi and Cirelli, 2003) proposed that the weight of synapses not important for learning are decreased. According to SHY, deep sleep and delta waves are involved in synaptic downscaling, which is likely to be reflected by a decrease of the firing rate. When considering our new definition of NREM substages, this would imply that N3, consisting in bursts of delta waves, should be associated with a decrease in global neuronal activity and that this decrease should persist even after the end of the N3 period. In order to take the variability across substages into account, we quantified the firing rate within NREM episodes when N3 epochs are preceded and followed by N2 periods, forming N2-N3-N2 triplets (Fig.6A). This analysis showed that the averaged firing rate decreased within N3, and that N2 firing rate in epochs following N3 is smaller on average than in epochs preceding N3, consistent with SHY. Nevertheless, the same is also true for N2 stage since the firing rate decreases within N2 (N3-N2-N3 triplets, Fig.6B). Altogether, this suggests that there is a global decrease in firing rate both during N2 and N3 but that the density of delta waves does not fully explain the long-lasting decrease of firing rate that is observed. Interestingly, the overall decrease in firing rate observed in N2 and N3 seemed to be compensated by a strong increase during and following REM sleep (Fig.6C). This could explain the global stability of the firing rate at the population level across sleep.

**Fig. 6.**
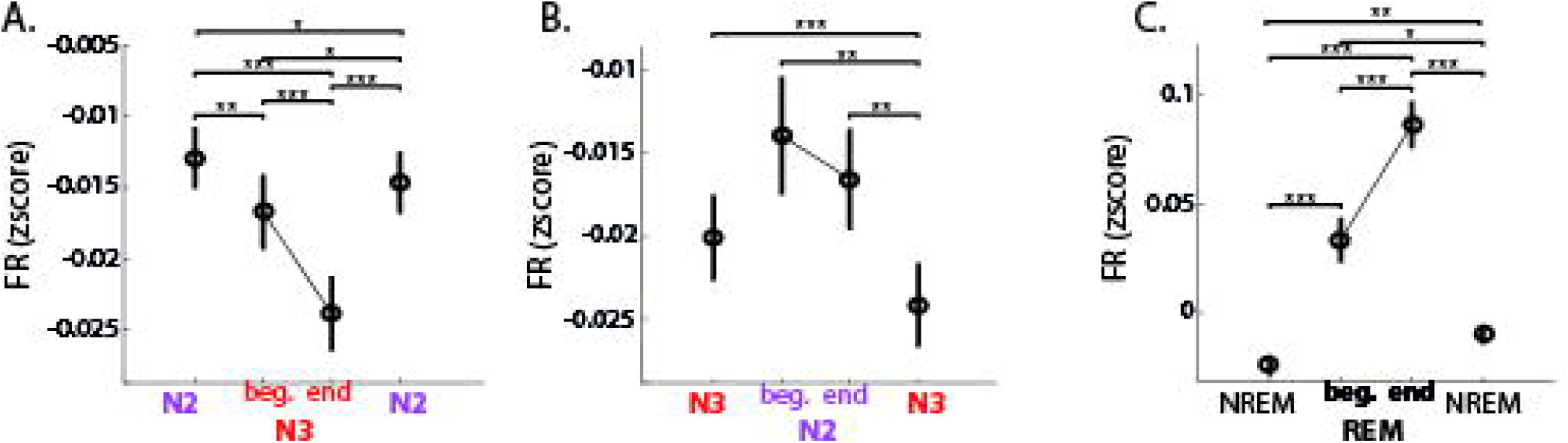
Firing of prefrontal neurons across substages. **(A-C)-** Normalized neuronal activity (zscore, n=591) averaged across all similar trio of epochs, for either N2-N3-N2 (A), N3-N2-N3 (B), or NREM-REM-NREM trio (C). Each episode is normalized in time to allow subsequent averaging, the 4 values correspond to the whole first epoch, the beginning (beg.) and the end of the second epoch, and the whole third one. *p<0.05, ** p<0.01, *** p<0.001, NS or not shown when not significant.

### NREM substages and the evolution of firing rate during sleep

Altogether, these results do not provide evidence for a selective role for bursts of delta waves as a mechanism to induce a long-lasting decrease of neuronal firing rate that is expected to be observed in case of synaptic depression predicted by SHY. Several other studies tested SHY hypothesis by looking at the evolution of firing rate across sleep but contradictory results have been observed (Vyazovskiy *et al*., 2009; Hengen *et al*., 2016; Watson *et al*., 2016; Cirelli, 2017). Here, we investigated whether taking NREM substages into consideration could improve our understanding of the neuronal activity evolution during sleep.

We thus quantified the evolution of the neuronal activity in mPFC during sleep. Firing rate were normalized (zscore) in order to avoid the high variability of the absolute firing rate (from 0.1Hz to 50Hz, Supplementary Fig.7A-B). In our data set, mPFC neurons were, on average, most active during REM sleep, their firing rate then decreasing from wakefulness, N1, N2 and finally N3 (Supplementary Fig.7A-D). It is worth pointing out that the difference of mPFC firing rate during N2 and N3 was due to the increase in down-states density, since firing rate in up-states is similar in both substages (Supplementary Fig.7B,D). Moreover, the correlation coefficient of neuronal activity between each pair of vigilance states reveals a strong difference between theses stages even at the neuronal level. Neurons that are more active in N2 are also active in N3 stages, while a negative correlation is found between wakefulness and the other sleep stages (Supplementary Fig.7G).

We then looked at the evolution of the normalized firing rate over the whole recording period. We did not observe any modification at the population level during sleep whatever the substage considered (Supplementary Fig.7E). A previous study found that the firing rate during wakefulness predicts the evolution of the firing rate during subsequent sleep: the high firing pyramidal neurons decreased their firing rate during sleep while the opposite was found for the low firing neurons (Watson *et al*., 2016). We found the same tendency with a significant positive correlation with time for the low firing pyramidal neurons, even if the negative correlation for the high firing neurons did not reach statistical significance in our data set (Supplementary Fig.8 and Supplementary Fig.9A-B).

We then addressed whether substages of NREM sleep could instead be used as predictors for the evolution of neuronal activity during sleep. As noticed in Fig.5A (and Supplementary Fig.5), transition between sleep substages seem to affect specific subpopulations of neurons. It is possible that the different NREM substages rely on a different balance of sleep-related neuromodulators, which could have different effect on neuronal firing rate. We thus differentiated five subpopulations of mPFC neurons, according to the sleep substage in which their firing rate is maximal (Fig.7A, see Methods for more detail). When averaging neuronal response of each preferring neuronal types at transition from one stage to another, we show that the modification of neuronal activity took place in a short time around transitions, supporting the fact that NREM substages reflects reliable changes in neuronal activity (Supplementary Fig.10). Notably, the transition to N1 from either wakefulness or REM is associated with an increase of N1-prefering neurons activity with a corresponding decrease of respectively the wake-preferring or the REM-preferring neurons.

**Fig. 7.**
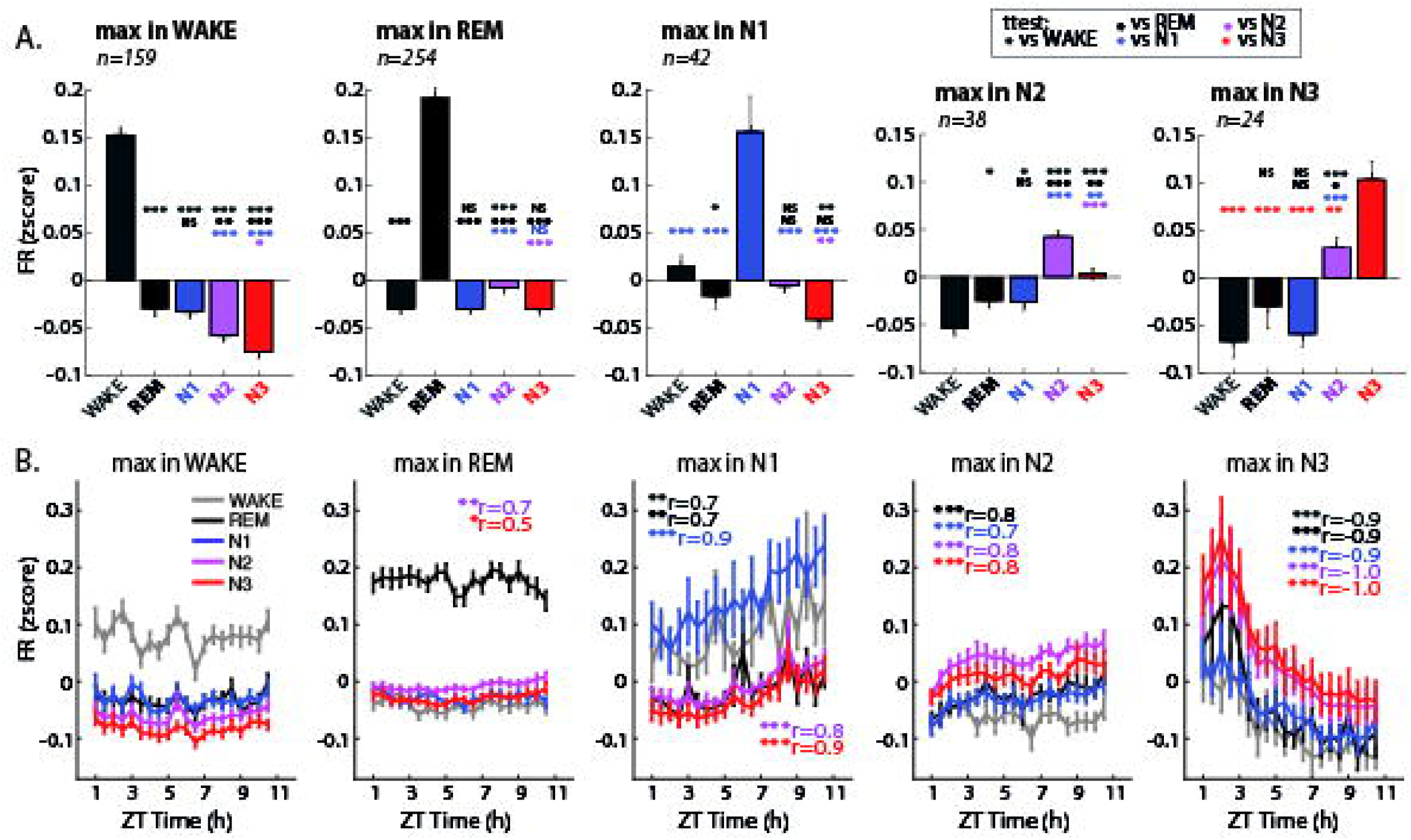
Evolution of cortical neurons activity depends on their “preferred” stage. **(A)-** Averaged normalized firing rate (zscore) of prefrontal cortical neurons restricted to each vigilance states, for all neurons subgroups based on their “preferred stage”: WAKE neurons fire more during wakefulness than other states, etc. **(B)-** Evolution across the light phase period of normalized firing rate during wakefulness and sleep stages for the different neuron subgroups. Spearman correlation coefficients are indicated for each substage in corresponding color. Note that to compare to the averaged normalized activity in panel A, the repartition of stages across the sleep period as shown in Fig.2 needs to be taken in consideration. * p<0.05, ** p<0.01, *** p<0.001, not significant are not shown.

The evolution profile across the whole sleep period was totally different across these neuronal groups: Wake-preferring and REM-preferring neurons, which represent around 75% of the data set neurons, did not change their activity across the sleep period. However, N1-preferring and N2-preferring neurons increased significantly their firing rate across the sleep period, while the N3-preferring neurons decreased strikingly their firing rate (Fig.7B, and Supplementary Fig.11D). These results are not due to the respective preponderance of each substages at the beginning and the end of the whole sleep session (Supplementary Fig.9C-D). The same specific evolution across time was found for pyramidal neurons (Supplementary Fig.11A-C). Importantly, these modifications were visible in all sleep stages, suggesting that the sleep and sleep-related electrophysiological events such as delta waves do not have a global effect on all cortical neurons and pointing out the existence of different population of neurons that should be taken into consideration. Altogether, these results support the existence of different neuronal subpopulations that are differently affected by sleep and that taking into account the existence of different substages of NREM sleep greatly enhance our understanding of the regulation of neuronal activity during sleep.

## Discussion

We present here a new sleep scoring method in mice differentiating three substages of NREM sleep inspired from human sleep scoring. We showed that most of the characteristics of human NREM substages were also found in mice sleep at the timescales of sleep period, sleep cycles, and stage transitions. Notably, the global sleep architecture, with the repartition of stages within sleep cycles being the reflection of the repartition observed at the whole sleep scale (Pittman-Polletta *et al*., 2013), is surprisingly well conserved between mice and human.

In all stages, we found striking similarities between rodent and human data. At the timescale of the whole sleep period, we observed an early high incidence of N3 followed by a progressive decline (Fig.2). Given that N3 identification is based on the presence of slow waves, this latter result is consistent with the well-described homeostatic regulation of sleep and the observation that slow waves can be used as a marker of sleep pressure (Tobler and Borbély, 1986; Huber *et al*., 2000; Vyazovskiy *et al*., 2009). Conjointly, we observed a progressive increase in the incidence of REM sleep, as reported in human sleep (Dement and Kleitman, 1957; Huber *et al*., 2000). Even more remarkably, these similarities still held when looking within a given sleep cycle. As in humans, REM sleep in mice occurred regularly defining sleep cycles that last around 10 min, compared to 90min in humans. In both species, cycles tend to start with N1, followed by N2, N3 and finally REM sleep. Finally, the transitions between sleep substages were also well conserved.

The main difference between mice and humans sleep stages concerns the duration of individual epochs which are one order of magnitude shorter in mice. This is consistent with theories suggesting that the duration of a sleep cycle and how well sleep is consolidated depend on a “homeostatic time constant”, which positively correlates with brain size and putative inertia of neuronal networks (Phillips *et al*., 2010). Accordingly, the dynamic of sleep in both human and rodent was predicted by a single biologically-inspired computational model, by only changing two variables: the homeostatic time constant and the mean inhibitory drive to the ventral-lateral preoptic nucleus (VLPO) (Phillips *et al*., 2010). This reproduced the shortness of sleep episodes and the high rate of transition observed in mice.

The shortness of the NREM substages episodes may explain why they have been neglected in mice studies so far. Several previous investigations already mentioned different activity profiles along NREM sleep in cats or rodents (Bergmann *et al*., 1987; Gottesmann, 1992; Neckelmann and Ursin, 1993; Rector *et al*., 2009), but without reaching the point of making a direct analogy with human NREM substages. Our study, by identifying a striking similarity between human and rodent sleep substages at the different timescales, suggests that although shorter the different sleep substages are conserved in both species.

Perhaps the most surprising observation was to find a sleep stage similar to human N1 in rodents. The observation of a sleep period with low amplitude activity has been reported several times in rodents and has been called either “sleep with low voltage EEG”, “low activity microstates” or “small-amplitude irregular activity” but has never been given a clear and operational definition (Bergmann *et al*., 1987; Lena *et al*., 2004; Miyawaki *et al*., 2017). Here, we showed that these periods do not occur in a random fashion but instead appear mostly at the beginning of the sleep cycle, a property also found in human sleep. This stage can also be differentiated from the other NREM substages at the neuronal level in the mPFC since there is no correlation between N1 and the other stages of sleep in the firing rate of the whole neuronal population. Moreover, the neuronal activity at transition from N1 and wakefulness highlights the difference between those stages (Supplementary Fig.10). The identification in rodent of a stage sharing homologies with the stage N1 in human has strong implication for the study of sleep pathologies and paves the way of new investigations. Indeed, N1 is increased in almost all disorders with sleep fragmentation such as insomnia or narcolepsy (American Psychiatric Association, 2013; Roth *et al*., 2013). We found the same increase in N1 in a mouse model of narcolepsy with a highly fragmented sleep.

In addition to the demonstration of a strong similarity between human and mice NREM substages, we also provided evidence supporting the notion that considering these substages greatly improves our understanding of the neuronal mechanisms at stake during sleep in relation to memory consolidation and homeostatic synaptic scaling.

Two theories about sleep and memory processes have been proposed. The reactivation theory posits that synaptic strength is increased by sleep reactivation during sharp waves ripples supporting memory consolidation. On the contrary, the second theory called SHY proposed that synaptic strength is decreased during sleep by a process relying on sleep slow/delta waves. In an attempt to reconcile the two theory, Genzel and colleagues proposed that consolidation would occur during light sleep and synaptic downscaling during deep sleep (Genzel *et al*., 2014), the latter statement in accordance with SHY (Tononi and Cirelli, 2003, 2014b). Yet the absence of a clear definition of light and deep sleep in rodent precludes a direct test of this model. Using our new NREM substages scoring, we observed a decrease in mPFC neuronal firing rate within single episodes of either N2 or N3. However, we showed that although there is a long lasting decrease of firing rate after N3 -a stage with high delta waves density, the same is also true after N2. These results do not support the hypothesis positing slow waves as a major effector of down scaling.

The evolution of firing rate during sleep has been previously investigated in order to test the SHY hypothesis, yet contradictory results were found (Vyazovskiy *et al*., 2009; Hengen *et al*., 2016; Watson *et al*., 2016; Cirelli, 2017). In a previous study, Vyazovskiy and colleagues (2009) found that the average firing rate of cortical neurons decreased over sleep following the natural evolution of sleep pressure consistently with the SHY hypothesis. However two other studies found that the evolution during sleep depends on the firing rate during wakefulness: high-firing pyramidal neurons decreases their activity over sleep whereas low-firing neurons activity increases, in line with a renormalization of firing rate during sleep (Grosmark *et al*., 2012; Watson *et al*., 2016). In our data set, we did not observe any global decrease of firing rate during sleep. We observed the same tendency as in Watson regarding the influence of the neuronal firing rate in the previous wake episodes on their evolution during subsequent sleep, yet it did not reach statistical significance for the low fining neurons.

However, we found that taking NREM substages into consideration could lead to a better prediction of the evolution of the firing rate during sleep. By looking at transition between sleep substages, it was possible to find neuronal populations that were differently modulated. This is consistent with the hypothesis that the different substages of sleep could reflect the changing neuromodulatory tonus that are known to evolve during sleep. There is indeed a large literature showing that variations in acetylcholine, noradrenalin and serotonin concentrations across sleep stages (Hasselmo, 1999) are associated with a modification of firing rate within selective population of neurons in subcortical structures (McGinty and Szymusiak, 2005). Distinct neuronal populations can respond differently to neuromodulators, as shown for instance by noradrenaline and acetylcholine response in the prefrontal cortex (Nagy *et al*., 2014; Uematsu *et al*., 2017). We thus classified the mPFC neurons regarding the substages in which their activity was maximal. We found that neurons firing preferentially during wakefulness or REM sleep did not change their activity during sleep. However, the N1-preferring and N2-preferring neurons increased their activity during sleep, while the neurons selective for N3 decreased their activity.

This shows that the evolution of the neuronal activity can be better predicted by the preferred sleep stage than the other factors considered so far. More generally, our results show that considering the existence of different substages of NREM sleep in rodents will greatly enhance our understanding of the regulation of neuronal activity during sleep.

As a conclusion, the strong similarity of sleep macro and microarchitecture between human and mouse we observed in our study legitimates the use of the rodent model to study sleep physiology and pathological states. We believe that the dysfunctions associated with deep slow wave sleep N3 such as night terrors or somnambulism (Kales *et al*., 1966; Fisher *et al*., 1973), and with N1 such as hypnagogic imagery, fibromyalgia (Wu *et al*., 2017), narcolepsy or persistent primary insomnia (Merica and Gaillard, 1992), will benefit from this new tool which will allow to address the complexity of those processes. Given the importance and the systematic use of stages classification in humans, the finding that the same scoring could be used in mice will pave the way for future studies on mechanisms that were neglected so far.

## Supporting information

## Acknowledgments

M.M.L. did the experiments and analyzed the data. J.L. did the experiments and analysis on Orexin-/-mice, provided by C.P.; G.d.L., S.B. and K.E. did several experiments included in the dataset. M.M.L. and K.B. wrote the manuscript, with helpful comment from Y.D., C.P. and S.L. Y.D. and S.L. provided expertise review on human analysis.

## Funding

This work was supported by the Fondation pour la Recherche sur le Cerveau (FRC), by the French National Agency for Research ANR- 12-BSV4-0013-02 (AstroSleep) and ANR-16-CE37-0001 (Cocode), by the CNRS: ATIP-Avenir (2014) and by the city of Paris (Grant Emergence 2014). This work also received support under the program Investissements d’Avenir launched by the French Government and implemented by the ANR, with the references: ANR-10-LABX-54 MEMO LIFE and ANR-11-IDEX-0001-02 PSL* Research University. M.M.L. and G.d.L. were funded by the Ministère de l’Enseignement Supérieur et de la Recherche, France.

## Competing interests

The authors report no competing interests.

