## Supplementary material for "Improved sleep scoring in mice reveals human-like stages"

### List of Supplementary Materials

Supplementary Methods

Supplementary Fig. 1. Schematic representation of the scoring method for NREM substages.

Supplementary Fig. 2. Frequency spectrograms in prefrontal cortex and hippocampus at stage transitions.

Supplementary Fig. 3. Evolution of Delta waves across sleep period and substages.

Supplementary Fig. 4. Substages repartition in early and late sleep cycles.

Supplementary Fig. 5. Mice sleep stage N1 characteristics in terms of movements and EMG.

Supplementary Fig. 6. Firing rates of prefrontal neurons averaged at stage transitions.

Supplementary Fig. 7. Firing rates of prefrontal neurons: normalization and evolution across sleep period.

Supplementary Fig. 8. Pyramidal versus interneurons classification and their firing characteristics.

Supplementary Fig. 9. Evolution of neuronal activity across sleep period for pyramidal neurons with high or low firing rate, interneurons, and simulated neurons.

Supplementary Fig. 10. Neuronal firing rates at stage transition, depending on neuron types.

Supplementary Fig. 11. Evolution of neuronal activity across sleep period depending on “preferred stage” for pyramidal neurons, and all neurons.

Supplementary Table. 1. DREAM Database of Human sleep recording sessions.

Supplementary Table. 2. Transitions between stages in mice and humans.

Supplementary Methods

Details on electrophysiological recordings and analysis

Signals from all electrodes were sampled at 20 kHz and digitalized by a headstage amplifier board equipped with accelerometer and then fed to a USB board connected to the acquisition computer (RHD2000-series, INTAN). Local field potentials were sampled and stored at 1,250 Hz. Electrophysiological signal analyses were performed with custom-made Matlab programs, based on generic code that can be downloaded at http://www.battaglia.nl/computing/ and <http://fmatoolbox.sourceforge.net/>.

Details on spike sorting and classification.

Tetrode recordings were bandpass-filtered between 600 and 9,000 Hz, then visualized and processed using NeuroScope and NDManager (http://neurosuite.sourceforge.net/). Whenever any of the tetrode channels exceeded a preset threshold, 1.6-ms samples were time-stamped and stored for all channels. Extracted waveforms were sorted via a semiautomatic cluster cutting procedure using Klusters (1) and KlustaKwik (2) (http://klustakwik.sourceforge.net/, now http://klusta.readthedocs.io/en/latest/), a program for unsupervised classification of multidimensional continuous data.

Method of classification between interneurons and pyramidal neurons included the calculation of firing rate, autocorrelogram and spike waveform parameters such as half amplitude width and trough to earlier or later peak *(3)*. The hyperplain dividing the two physiologically identified classes was used to separate units into putative interneurons and putative pyramidal cells.

Details on human recordings

We used the DREAMS database, which is entirely attributed to University of MONS - TCTS Laboratory (Stéphanie Devuyst, Thierry Dutoit) and Université Libre de Bruxelles - CHU de Charleroi Sleep Laboratory (Myriam Kerkhofs). Additional information can be found at http://www.tcts.fpms.ac.be/~devuyst/Databases/DatabaseSubjects. Those polysomnographic recordings correspond to 20 whole-nights from healthy subjects, 16 women and 4 men from 20 to 65 years old (mean 33,4), and are in EDF format (polygraphe BrainnetTM System of MEDATEC, Brussels, Belgium) sampled at 200Hz, including 2 electro- occulograms, 3 EEG (Fpl-A2, Cz-A1 and O1-A2) and one chin EMG. These recordings were selected for their clarity (i.e. that they contain few artifacts) and come from persons, free of any medication, volunteers in other research projects, conducted in the sleep lab. Sleep scoring was performed by an expert from the Sleep Laboratory, according to the new standard of the American Academy of Sleep Medicine *(4)*.

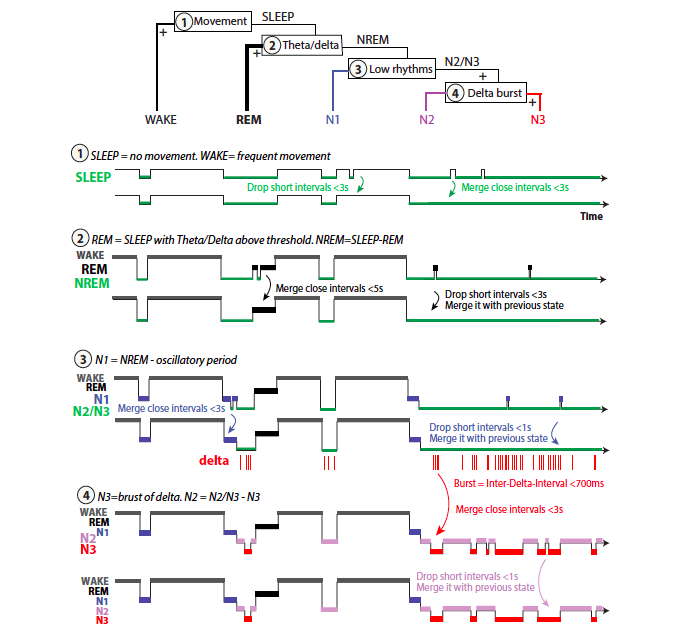

**Sup Fig. 1. Schematic representation of the scoring method for NREM substages.**

Steps 1 and 2 are inspired from classical sleep scoring, but merge and drops which is usually done by visual continuity have been automatized. Step 3 and 4 allow the sub-classification of NREM into N1, N2 and N3 based on oscillatory activity in the low frequency bands (2-20Hz) and delta waves in the prefrontal cortex.

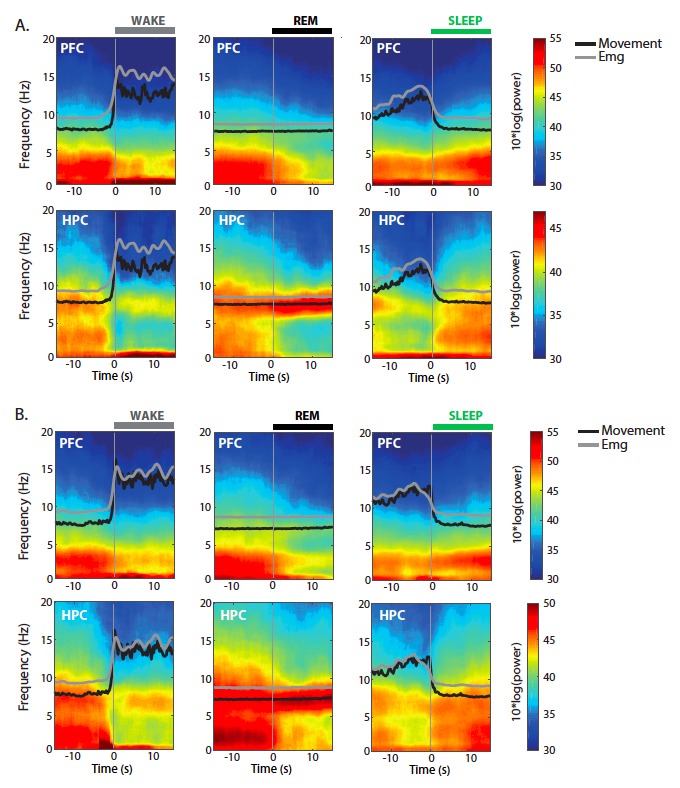

**Sup Fig. 2. Frequency spectrograms in prefrontal cortex and hippocampus at stage transitions.**

**(A)-** Low frequency (0-20Hz) spectrogram in medial prefrontal cortex (PFC) and hippocampus (HPC) averaged on all transitions to wakefulness (left, n=45 transitions), to REM (middle, n=79 transitions) and to sleep before any substage scoring (right, n=108 transitions) shown in a 30s window (N=1 mouse sleep session). Averaged EMG signal (grey) and movement quantity (black) are also shown in superposition after normalization (zscore). **(B)-** Same as (A) for one different mouse (N=1). Spectrogram in PFC and HPC averaged on all transitions to wakefulness (left, n=64 transitions), to REM (middle, n=60 transitions) and to sleep before any substage scoring (right, n=107 transitions).

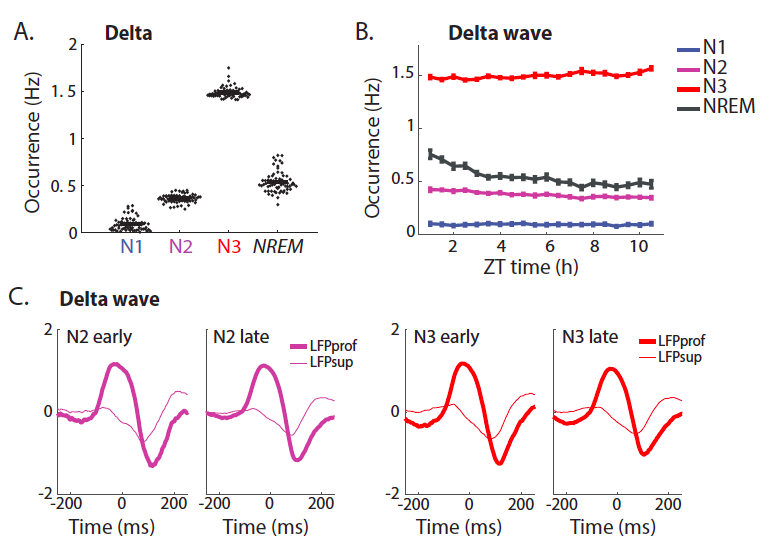

**Sup Fig. 3. Evolution of Delta waves across sleep period and substages.**

**(A)-** Occurrence of prefrontal delta waves averaged on each stages periods N1, N2, N3, or all NREM for comparison. Those differences reflect the criteria of NREM sleep scoring. Each point represents one night, bars represent the mean, n=65, N=25. **(B)-** Evolution across sleep period of delta wave occurrence in mice. Rhythms are restricted either to N1, N2, N3 or all NREM and averaged on successive 30min periods across the sleep period. Note the decrease of delta wave density during NREM but no evolution within each substage. N=15, error bars show SEM. **(C)-** Comparison of averaged delta wave shape during N2 (left) or N3 (right) between the beginning and the end of the sleep period (n=1 sleep session).

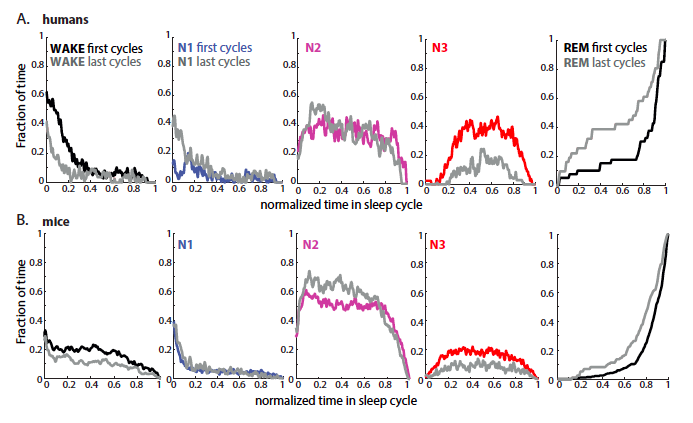

**Sup Fig. 4. Substages repartition in early and late sleep cycles.**

**(A-B)-** Comparison between humans (A, N=20 subjects) and mice (B, n=33 sleep sessions, N=13 mice) sleep cycle content in terms of vigilance states, either at the beginning of sleep period (colored) or at the end (grey). All cycles terminate by REM sleep by definition, only long enough cycles were taken in consideration (15s and longer for mice, 30min and longer for humans). Note that cycles are richer in REM and poorer in N3 in the last cycles for both humans and mice.

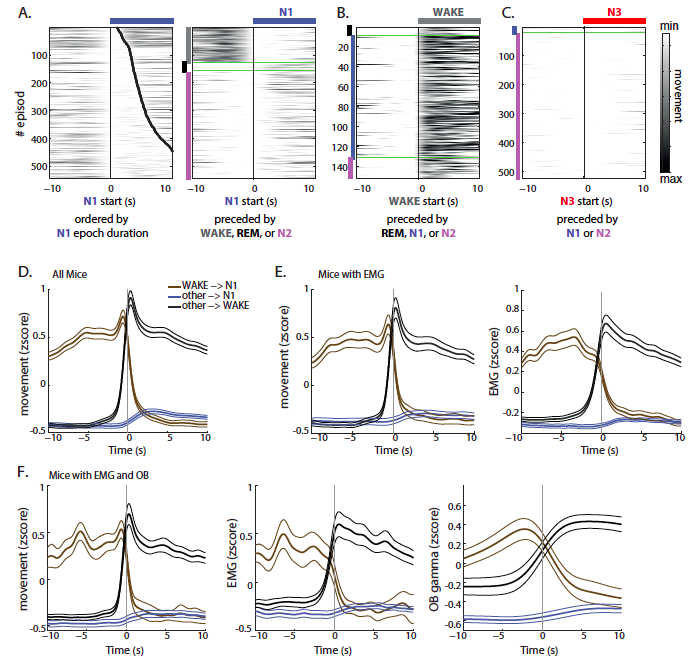

**Sup Fig. 5. Mice sleep stage N1 characteristics in terms of movements and EMG.**

**(A)-** Example of movements quantity (arbitrary units, see Methods) averaged on 20s period aligned on the beginning of N1 episode. Row are ordered by the duration of N1 episode, black dots indicating the end of episode, or by the nature of the preceding episode: Wake, REM or N2 (indicated by grey, black or pink lines, respectively). Note that N1 is not associated with movement, which could indicate that electrophysiological signal change is due to short awakening. **(B)-** Movements quantity for the same example averaged on 20s period aligned on the beginning of wakefulness episode, ordered by the nature of the preceding episode: REM, N1 or N2. **(C)-** Same as (B) aligned on the beginning of N3 episodes, ordered by the nature of the preceding episode: N1 or N2. Some transitions are not represented due to their rarity. **(D)-** Quantity of movement, normalized (zscore) on individual recording, averaged on transition from wakefulness to N1, from any sleep stage to N1, or from any sleep stage to Wakefulness, for all mice recording sessions (n=56, N=22). **(E)-** Normalized (zscore) quantity of movement and EMG, averaged on same transitions as in (D), for mice with additional EMG recording (n=29, N=10). **(F)**- Normalized (zscore) quantity of movement, EMG and high gamma power (averaged on 50-70Hz band) recorded in the olfactory bulb (OB), averaged on same transitions as in (D), for mice with additional EMG and OB recording (n=19, N=7).

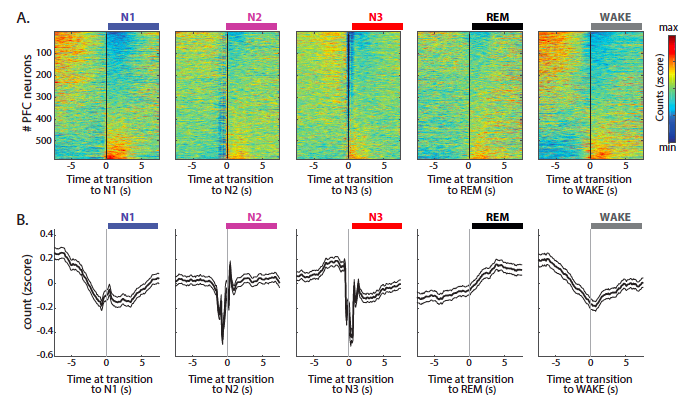

**Sup Fig. 6. Firing rates of prefrontal neurons averaged at stage transitions.**

**(A)-** Peri-event histograms of prefrontal cortex neurons (n=591 neurons) activity at transition to N1, N2, N3, REM and wakefulness, ordered by the intensity of response to transitions. Each row corresponds to the cross-correlogram of one neuron spikes by all transitions to indicated stage, normalized by zscore. **(B)-** Quantification of peri-event histograms presented in (A), averaged on all neurons. Note the sharp neuronal response to all stages transitions and the drop of neuronal activity just before N2 start and at the beginning of N3 corresponding to down states.

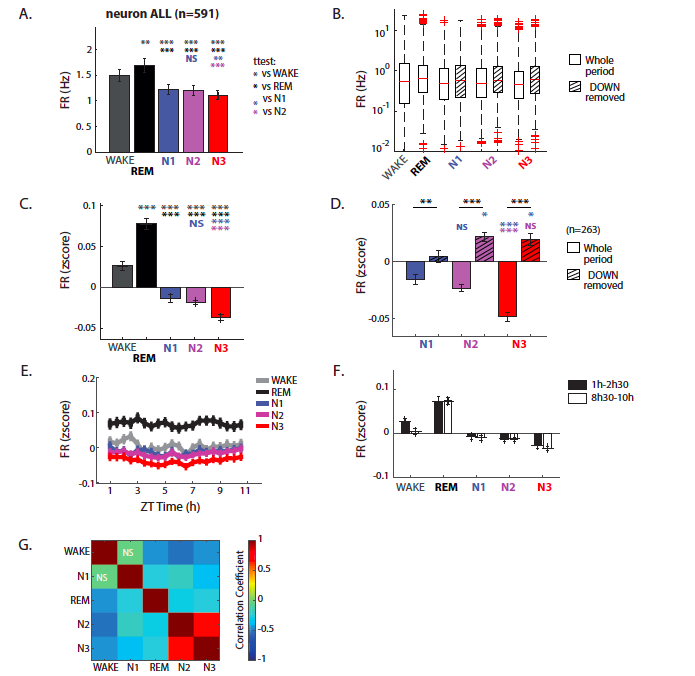

**Sup Fig. 7. Firing rates of prefrontal neurons: normalization and evolution across sleep period.**

**(A)-** Averaged firing rate across vigilance states: wakefulness, REM sleep, and NREM substages N1, N2, N3. **(B)-** Box plot for firing rate across vigilance states: wakefulness, REM sleep, and NREM substages N1, N2, N3. Dashed box represent the firing rate measured on periods outside the detected down states. **(C-D)-** Averaged firing rate of neurons after normalization (zscore). For each neuron (n=591), instantaneous firing rate of the whole recording is normalized (zscore), then averaged on each stage (C). Dashed bars indicate firing rate averaged on stages after down states periods have been removed (D). As expected normalized firing rate is increased after removing down state, although this difference is significant only in N3. **(E-F)-** Evolution of normalized neuronal activity (zscore, n=591). Evolution of normalized firing rate across sleep period, during wakefulness and sleep stages (E) and comparison between the first 90min and last 90min of the sleep period (F), for all prefrontal neurons (n=591). **(G)-** Matrix of correlation between sleep stages for normalized neuronal activity. NS for no significant correlation.

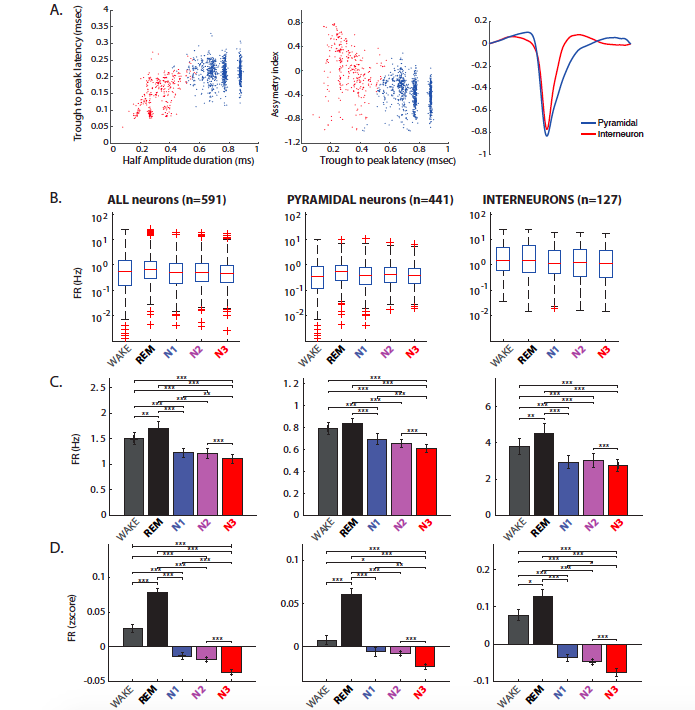

**Sup Fig. 8. Pyramidal versus interneurons classification and their firing characteristics. (A)-** Classification between pyramidal (blue) and interneurons (red) based on waveforms characteristics. With method of classification based on Sirota and colleagues work *(3)*, neurons were clustered according to waveform asymmetry and mean filtered spike width. Putative excitatory and inhibitory neurons form separate clusters and have different mean waveforms (right panel). **(B-D)-** Box plot for firing rate (B), averaged firing rates (C) and averaged normalized (zscore) firing rates (D) across vigilance states: wakefulness, REM sleep, and NREM substages N1, N2, N3. * p<0.05, ** p<0.01, *** p<0.001, not significant are not shown.

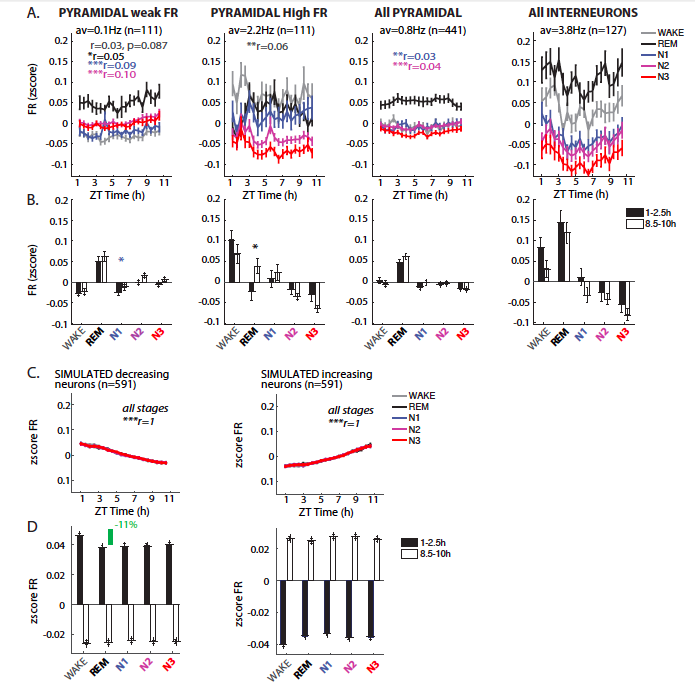

**Sup Fig. 9. Evolution of neuronal activity across sleep period for pyramidal neurons with high or low firing rate, interneurons, and simulated neurons.**

**(A)-** Evolution of neuronal activity across sleep period for pyramidal neurons, either presenting weak firing rate (mean=0.1Hz) or high firing rate (mean=2.8Hz) as in *(5)*, all pyramidal neurons and all interneurons (from left to right). Correlation coefficient of Spearman test are indicated for each substage in the corresponding color. Not significant correlations are not shown. **(B)-** Comparison of normalized neuronal activity (zscore) between the first and last hours of the sleep period for neurons group as in (A). **(C)-** Evolution of neuronal activity across light phase for simulated neurons with regularly increasing (left) or decreasing (right) firing rate across the sleep period. Simulated neurons (n=591) have the same repartition of firing rates as in the recorded data, firing rate is normalized (zscore) across the whole period and activity is then averaged on each substage for each neuron. **(D)-** Comparison of normalized neuronal activity (zscore) between the first 90min and last 90min of the sleep period for neurons group as in (C). Note that in our simulated neurons, the maximal difference between stages reach 11%, suggesting that the difference observed in our real data from one substage to the other cannot be explained by random or continuous evolution of firing rates across sleep period.

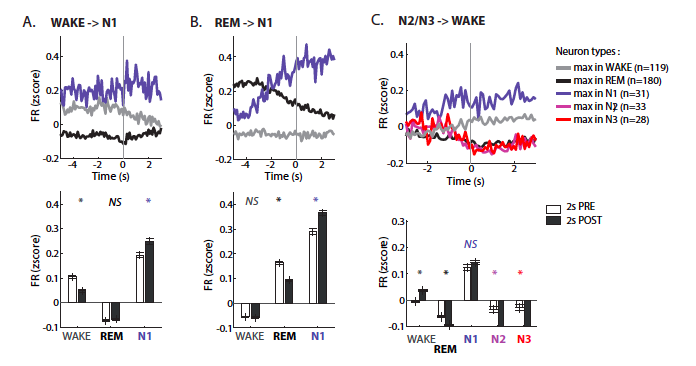

**Sup Fig. 10. Neuronal firing rates at stage transition, depending on neuron types.**

**(A-B)-** Averaged normalized firing rates (zscore) for WAKE-, REM- and N1-prefering prefrontal neurons at the transition from wakefulness to N1 (A) and from REM to N1 (B). Note the increase of N1 neurons activity at N1 onset, which is not accompanied by WAKE neurons activity increase. Bottom panels corresponds to quantification of the averaged normalized firing rates presented in the upper panels, averaged on the 2s before transition (white bars) and 2s after transition (grey bars) for comparison. **(C)-** Averaged normalized firing rates (zscore) for WAKE-, REM-, N1-, N2- and N3-prefering neurons at the transition from N2/N3 to wakefulness. Bottom panel corresponds to quantification of the averaged normalized firing rates presented in the upper panel, averaged on the 2s before transition (white bars) and 2s after transition (grey bars) for comparison. * p<0.05, ** p<0.01, *** p<0.001, NS for not significant.

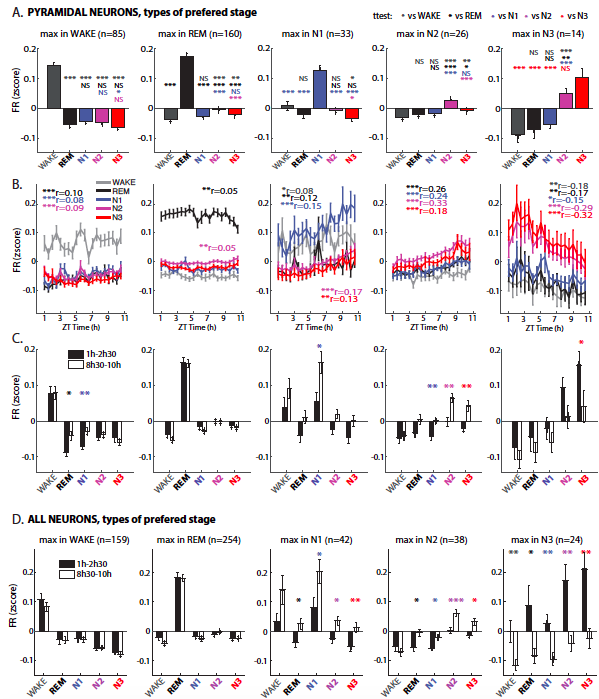

**Sup Fig. 11. Evolution of neuronal activity across sleep period depending on “preferred stage” for pyramidal neurons, and all neurons.**

**(A)-** Activity of prefrontal cortical pyramidal neurons based on their “preferred stage”: WAKE neurons fire the highest during wakefulness, REM neurons during REM sleep, etc. Averaged firing rate of neurons after normalization (zscore) restricted to all 5 vigilance states. **(B)-** Evolution of normalized firing rate across sleep period during wakefulness and sleep stages for the different subgroups of pyramidal neurons. Correlation coefficient of Spearman test are indicated for each substage in the corresponding color. **(C)-** Comparison of normalized neuronal activity (zscore) between the first 90min and last 90min of the sleep period for all subgroups of pyramidal neurons. **(D)** Same quantification as in (C) for all prefrontal cortex neurons presented in Fig.8. * p<0.05, ** p<0.01, *** p<0.001, not significant are not shown.

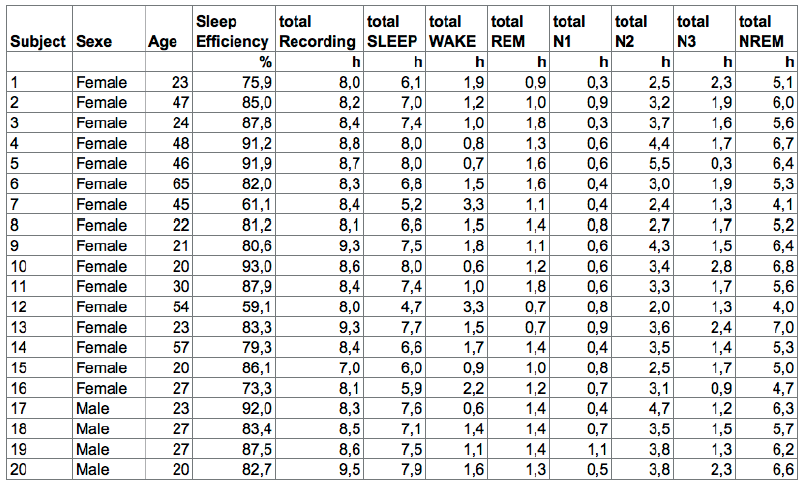

**Sup. Table. 1. DREAM Database of Human sleep recording sessions.**

Information on all sleep recording sessions from the 20 subjects of the DREAM database.

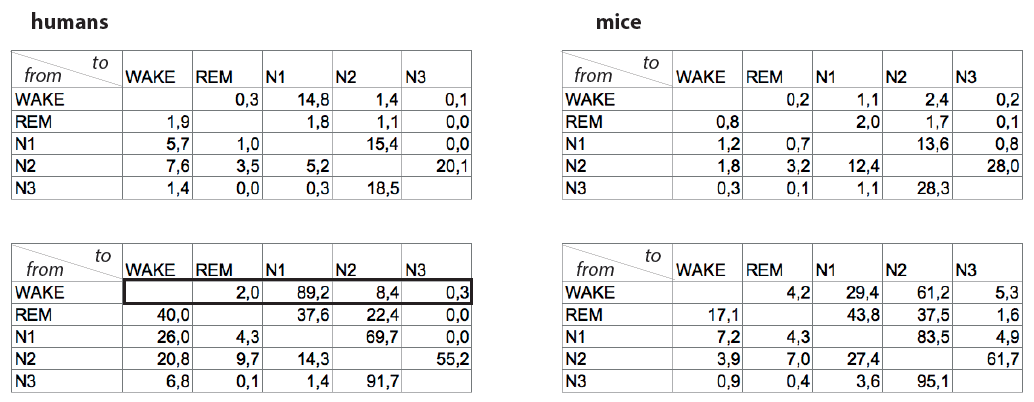

**Sup Table. 2. Transitions between stages in mice and humans.**

Averaged number of transitions from one stage (wakefulness, REM sleep, N1, N2 or N3) to another for all subjects (left) and all mice (right), either expressed as a percentage of all transitions recorded in the session (upper panels) or as a percentage of all transition from one stage (bottom panels). For example, the first row of the latter panel (thick lines) would be the transition from wakefulness to REM sleep, N1, N2 and N3 respectively, expressed as a percentage of all transition from wakefulness, the sum of each row therefore being 100.
